# FDA-approved drug screening identified micafungin as an antiviral agent against bat-borne emerging zoonotic Pteropine orthoreovirus

**DOI:** 10.1101/2022.09.21.508823

**Authors:** Tetsufumi Katta, Ayato Sato, Naoya Kadofusa, Tomoki Ishibashi, Hiroshi Shimoda, Atsuo Iida, Eiichi Hondo

**Affiliations:** Laboratory of Animal Morphology, Graduate School of Bioagricultural Sciences, Nagoya University, Nagoya 464-8601, Japan; Institute of Transformative Bio-Molecules (WPI-ITbM), Nagoya University, Nagoya 464-8601, Japan; Laboratory for Physical Biology, RIKEN Center for Biosystems Dynamics Research, Kobe 650-0047, Japan; Laboratory of Veterinary Microbiology, Joint Graduate School of Veterinary Medicine, Yamaguchi University, 1677-1 Yoshida, Yamaguchi, 753-8515, Japan

**Keywords:** Pteropine orthoreovirus, Antiviral, FDA-approved drug, Micafungin

## Abstract

Bat-borne emerging zoonotic viruses cause major outbreaks, such as the Ebola virus, Nipah virus, severe acute respiratory syndrome (SARS) coronavirus, and SARS-CoV-2. Pteropine orthoreovirus (PRV), which spillover event occurred from fruit bats to humans, causes respiratory syndrome in humans widely in South East Asia. Repurposing approved drugs against PRV is a critical tool to confront future PRV pandemics. We screened 2,943 compounds in an FDA-approved drug library and identified eight hit compounds that reduce viral cytopathic effects on cultured Vero cells. Real-time quantitative PCR analysis revealed that six of eight hit compounds significantly inhibited PRV replication. Among them, micafungin used clinically as an antifungal drug, displayed a prominent antiviral effect on PRV.

**Highlights:**

- A library of 2,943 FDA-approved drugs was screened to find potential antiviral drugs of Pteropine orthoreovirus.
- Six hit compounds dramatically inhibited viral replication *in vitro*.
- Micafungin possessed antiviral activity to multiple strains of PRV.

## 1. Introduction

Emerging infectious diseases that have emerged in recent years and caused outbreaks worldwide pose a major threat in today’s human society, where movement across borders and continents has become more accessible. The Ebola virus (Leroy et al., 2005) has repeatedly emerged mainly in West and Central Africa. Nipah virus (Chua et al., 2002) caused outbreaks in Malaysia and India. SARS-CoV (Li et al., 2005) and SARS-CoV-2 (Zhou et al., 2020) caused a global pandemic. All of these viruses are considered to be of bat origin. Bats do not show severe clinical symptoms when infected with these viruses, and they act as carriers of the virus by flying while infected (Middleton et al., 2007; Swanepoel et al., 1996; Watanabe et al., 2010). Bat-borne viruses have emerged due to indirect transmission through other wildlife or direct transmission from bats to humans (Irving et al., 2021; Wang and Anderson, 2019). Surveillance of bat viruses and the establishment of medical treatments are critical to prevent damage from infectious diseases.

Pteropine orthoreovirus (PRV) is one such bat-borne virus. PRV belongs to the genus *orthoreovirus*, family *Reoviridae*. The first isolation was from flying foxes in 1968 (as Nelson Bay orthoreovirus) (Gard and Compans, 1970). It was considered a virus that only bats have had for a long time. Still, in 2006, PRV was isolated from patients with acute respiratory symptoms, and its pathogenicity in humans was confirmed (Kaw et al., 2007). Until now, PRV has been isolated from flying foxes (Pritchard et al., 2006; Takemae et al., 2018; Taniguchi et al., 2017) and patients (Cheng et al., 2009; Chua et al., 2011; Wong et al., 2012; Yamanaka et al., 2014) in Malaysia, Indonesia, and Philippine. Most patients with PRV infection suffer from cough, sore throat, and high fever, and some develop symptoms of infectious gastroenteritis (diarrhea and vomiting) (Tan et al., 2017). A recent study reported that 13%-18% seropositivity against PRV remained from 2001 to 2017 on Tioman Island, Malaysia (Leong et al., 2022).

More attention should be paid to the genomic structure of PRV. The segmented RNA genome of PRV is the driving force for viral evolution by genetic reassortment (McDonald et al., 2016). Progeny virus resulting from genomic segment exchange during co-infection of various virus strains may have altered host fitness. In influenza virus with a similarly segmented RNA genome, genetic reassortment occurred in pigs, and the H1N1/2009 virus infecting humans caused a major pandemic (Smith et al., 2009). Possible genetic reassortment events among PRV strains have already been reported (Takemae et al., 2018). Therefore, PRV is an emerging infectious bat-borne virus that poses a future threat of altered pathogenicity, but no specific clinical treatment for PRV infection has yet been developed.

Drug repurposing is a practical approach to combat emerging infectious diseases. Officially approved medical drugs have well-recognized safety, require less time for approval in another usage, and are cost-effective to manufacture than new chemical drugs (Mercorelli et al., 2018). Thus, these drugs are potent in the early stages of an emerging infectious disease epidemic. During the SARS-CoV-2 pandemic, an inhibitor of the Ebola virus RNA-dependent RNA polymerase, Remdesivir (Veklury; Gilead Sciences, USA) used for the treatment (Wang et al., 2020). In this study, we performed in vitro screening using an FDA-approved drug library as a key for drug repurposing against PRV infections.

Fusogenic orthoreovirus to which PRV belongs does not have an envelope and induces rapid (approximately 8 hours post-infection) syncytium formation when they infect cultured cells via a specialized nonstructural protein FAST (Ciechonska and Duncan, 2014). Reduced lethality of FAST-deficient PRV in mice indicates that syncytium formation is an important step for virulence (Kanai et al., 2019). In this study, we used this inhibition of syncytial formation to evaluate the drugs.

## 2. Materials and methods

### 2.1 Cells

Vero JCRB 9013 cells were used in this study. Cells were cultured in Dulbecco’s Modified Eagle’s Medium(DMEM; Nissui, Tokyo, Japan) containing 10% fetal bovine serum (FBS; HyClone, Logan, USA), 2% L-glutamine (Sigma, Milwaukee, Wisconsin, USA), 0.14% sodium hydrogen carbonate (NaHCO_3_; Sigma, Milwaukee, USA), and 100 U/mL·0.1 μg/mL penicillin-streptomycin (Meiji, Tokyo, Japan).

### 2.2 PRVs

The viral strains Garut-50 (PRV50G) and Garut-69 (PRV69G) that were previously isolated from greater flying fox (*Pteropus vampyrus*) in Indonesia (Takemae et al., 2018) were propagated in Vero cells in DMEM medium containing 2% FBS and stored at −80°C until use. Virus titration was performed by plaque assay in Vero cells.

### 2.3 Chemical compounds

Of the 2,943 chemical compounds in the FDA-approved drug library (L1300, Selleck, Houston, USA), 2,594 compounds were pre-dissolved to 10 mM in DMSO, and 257 were in the water. Seventy-four were pre-dissolved to 2 mM in DMSO and 18 in water. The compounds were stored at −30°C until use. Gemcitabine (G0367, TCI, Tokyo, Japan), micafungin sodium (HY-16321, Medchemexpress, New Jersey, USA), and chlorophyllin sodium copper salt (C2945, LKT, Minnesota, USA) were additionally purchased.

### 2.4 Syncytium inhibition assay

Syncytium inhibition assay was performed to PRV50G. Vero cells were seeded at a concentration of 2 × 10^4^ cells/well in a 96-well plate, cultured in DMEM containing 2% FBS, and incubated for 24 h at 37°C, 5% CO2. Each compound of the FDA-approved drug library was added to each well at a concentration of 20 µM, and two hours later, PRV50G was infected at MOI = 0.1. The cytopathic effect (CPE) was observed by light microscopy. Compounds added to wells with less CPE than controls or no CPE observed were selected.

### 2.5 MTT assay

MTT assay was performed to measure the cell viability of PRV50G infected Vero cells. MTT solution of MTT Cell Count Kit (Nacalai tesque, Kyoto, Japan) was added to each well of section 2.4. After two hours of reaction, the solubilizing solution was added and incubated at 37°C overnight. The absorbance values at 595 nm and 750 nm were measured in a microplate reader. After correcting the absorbance value with blank, the relative cell viability was calculated as the ratio of the absorbance value of the mock group. Each experiment was triplicated.

### 2.6 RT-qPCR

For further validation of the anti-PRV50G effect of eight hit compounds and 3’,5’-Di-O-benzoyl-2’-deoxy-2’,2’-difluorocytidine, a prodrug of gemcitabine, qPCR was performed to quantify viral load in PRV50G infected Vero cells. Each of the nine compounds was diluted with DMSO to 2, 8, 20, and 50 µM. Each dilution or DMSO was added 1 µl to Vero cells cultured in a 96-well plate, and after two hours, PRV50G was infected at MOI = 0.01. At 24 hours post-infection (hpi), the supernatant was removed, cells were washed with PBS, and used as a sample for RNA extraction.

Total RNA was extracted using ISOGEN2 (Nippon gene, Tokyo, Japan). Reverse transcription reaction and qPCR reaction were performed in one step using RNA-direct SYBR Green Realtime PCR Master Mix (TOYOBO, Osaka, Japan) (90°C for 30 s, 61°C for 20 min, 95°C for 30 s, 95°C for 5 s −55°C for 10 s - 74°C for 15 s for 35 cycles). The melting curve was analyzed (95°C for 10 s, 65°C for 60 s, and 97°C for 1 s). Primer sequences were designed to amplify a region of the S4 segment region of PRV50G (forward,5’-TTGGATCGAATGGTGCTGCT-3’; reverse, 5′-TCGGGAGCAACACCTTTCTC-3’, amplicon size: 159 bp). A standard curve was created and obtained Ct values were plotted. The relative viral RNA copy number compared to the DMSO group was calculated. Each experiment was hexaplicated.

### 2.7 ATP assay

ATP assay was performed to confirm the cytotoxicity of the six hit compounds and the gemcitabine prodrug, except for evans blue and caspofungin, which had low PRV inhibitory effects among the hit compounds. Each of the seven compounds or DMSO was added 1 µl at concentrations of 2, 8, 20, and 50 µM to Vero cells cultured in a 96-well plate. After incubation at 37°C for 24 hours, CellTiter-Glo 2.0 Cell Viability Assay solution (Promega, USA) was added. Luminescence values were measured by Infinite 200 PRO (TECAN, Switzerland). The relative cell viability was calculated as the ratio of the luminescence value of the vehicle group. Each experiment was hexaplicated.

### 2.8 Verification of antiviral steps

Binding / Entry / Post-entry assays were performed to validate the antiviral steps of the six hit compounds to PRV50G infection. In the Binding assay, Vero cells cultured in 96-well plates as in section 2.4 were preincubated at 4°C for 30 min, PRV50G at MOI = 0.1, and each compound was added simultaneously for 90 min adsorption reaction at 4°C. In the Entry assay, PRV50G was inoculated at MOI=0.1 after preincubation at 4°C for 30 min, and the adsorption reaction was performed at 4°C for 90 min. After washing PBS twice, each drug was added and incubated at 37°C for 90 min. In the Post-entry assay, PRV50G was inoculated at MOI = 0.01, incubated at 37°C for 90 min, washed twice with PBS, and incubated for 24 h with each compound. In each treatment, RNA was extracted, and the relative viral copy number was calculated by RT-qPCR as in section 2.5. Each experiment was triplicated.

### 2.9 Compound effects on another strain of PRVs

To test whether micafungin inhibits another strain of PRVs at the post-entry step, we also examined its effect on PRV69G. Vero cells were seeded at a concentration of 1.5 × 10^5^ cells/well in a 24-well plate, cultured in DMEM containing 2% FBS, and incubated for 24 h at 37°C, 5% CO2. PRV50G or PRV69G were infected at MOI = 0.01 or 0.1, respectively, incubated at 37°C for 90 min, and washed twice with PBS. DMSO or micafungin was added to each well, incubated for 24 hours, and replaced with a 0.8% agarose-containing medium. After 24 or 48 hours, cells were stained with crystal violet. The plaque area that was not stained was measured by ImageJ Fiji. The threshold value was set at 20-180.

### 2.10 Statistics

All statistical analyses were performed using the statistical software R (ver. 4.2.0). Statistical differences were calculated by the Wilcoxon rank sum test or students’ T-test. The threshold value is P < 0.05.

## 3. Results

### 3.1 Identification of Hit compounds

To identify chemical compounds with anti-PRV activity, 2,943 compounds from the FDA-approved drug library were screened. First, primary hit compounds that inhibit the cytopathic effect (CPE) that PRV exhibits in Vero cells were identified (Fig. 1A). PRV induces syncytium formation when it infects cultured cells. The syncytium eventually dissolves as viral replication proceeds, leaving traces of the syncytium in the remaining dish (Fig. 1B). Compounds show smaller syncytium formation than that of the control, or no syncytium formation was observed were selected. As a result, 27 primary hit compounds were obtained. In addition, MTT assays were performed to measure the viability of infected cells (Fig. 1C). The top eight compounds with the highest cell viability were selected as hit compounds (Fig. 1D).

**Fig. 1.**
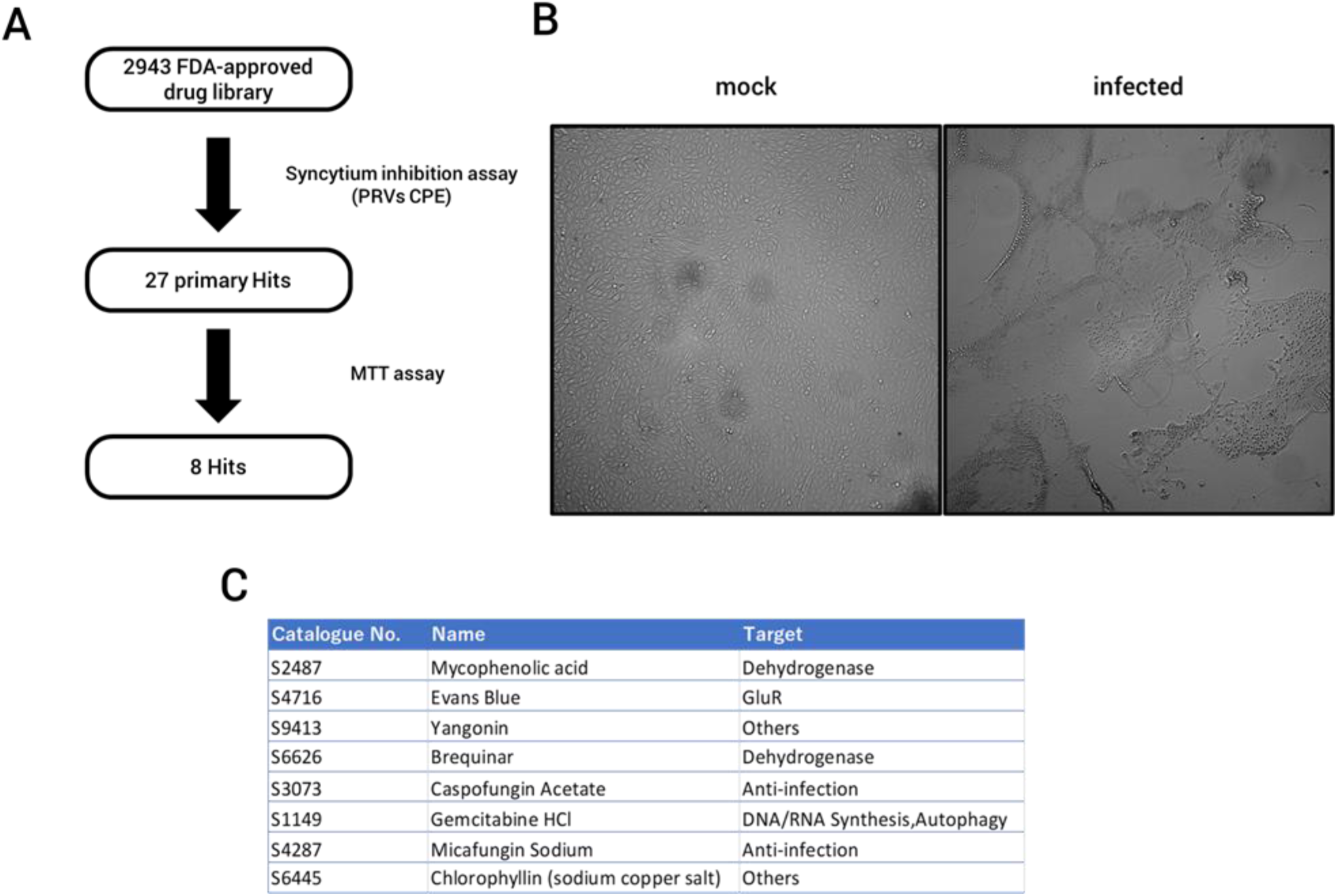
Phenotypic screening identified eight hit compounds. (A) Schematic representation of screening. (B) Cytopathic effect (CPE) of PRV50G. Vero cells infected with PRV50G at MOI = 0.1 for 24 h. The picture captured by light microscopy. (C) The relative cell viabilities of Vero cells infected with PRV50G. (n = 3, mean ± SE) The name of each compounds represented by catalogue number of the supplier. (D) List of eight hit compounds.

### 3.2 Inhibitory effects of hit compounds on PRV replication

The anti-PRV activity of hit compounds was verified by RT-qPCR. Vero cells which were treated with each of eight hit compounds or 3’,5’-Di-O-benzoyl-2’-deoxy-2’,2’-difluorocytidine, the gemcitabine prodrug, were infected PRV50G. Of the nine compounds, mycophenolic acid, yangonin, brequinar, gemcitabine, gemcitabine prodrug, micafungin, and chlorophyllin decreased relative viral plaque RNA copy number less than 1 × 10^−2^ at 50 µM (Fig.2A). The highest inhibition was observed in gemcitabine prodrug at 1 × 10^−5.81^. Mycophenolic acid, brequinar, and gemcitabine greatly inhibited PRV50G replication at a low concentration range (2 µM). Micafungin inhibited PRV50G replication in a dose-dependent manner, increasing its efficacy to levels comparable to gemcitabine (1 × 10^−4^) at 50 µM. Secondary, cell cytotoxicity of seven compounds, which exhibited high inhibition to PRV50G by RT-qPCR, was measured by ATP assay (Fig. 2B). Unlike the MTT assay, the ATP assay can eliminate the reducing properties of the compound. Without virus infection, the amount of ATP was measured 24 hours after the addition of the compound, and the relative cell viability compared to the control group was calculated. The results showed that cell viability did not decrease below 75% up to 50 µM for the six compounds, mycophenolic acid, yangonin, brequinar, gemcitabine, micafungin, chlorophyllin, except gemcitabine prodrug (Fig. 2B).

**Fig. 2.**
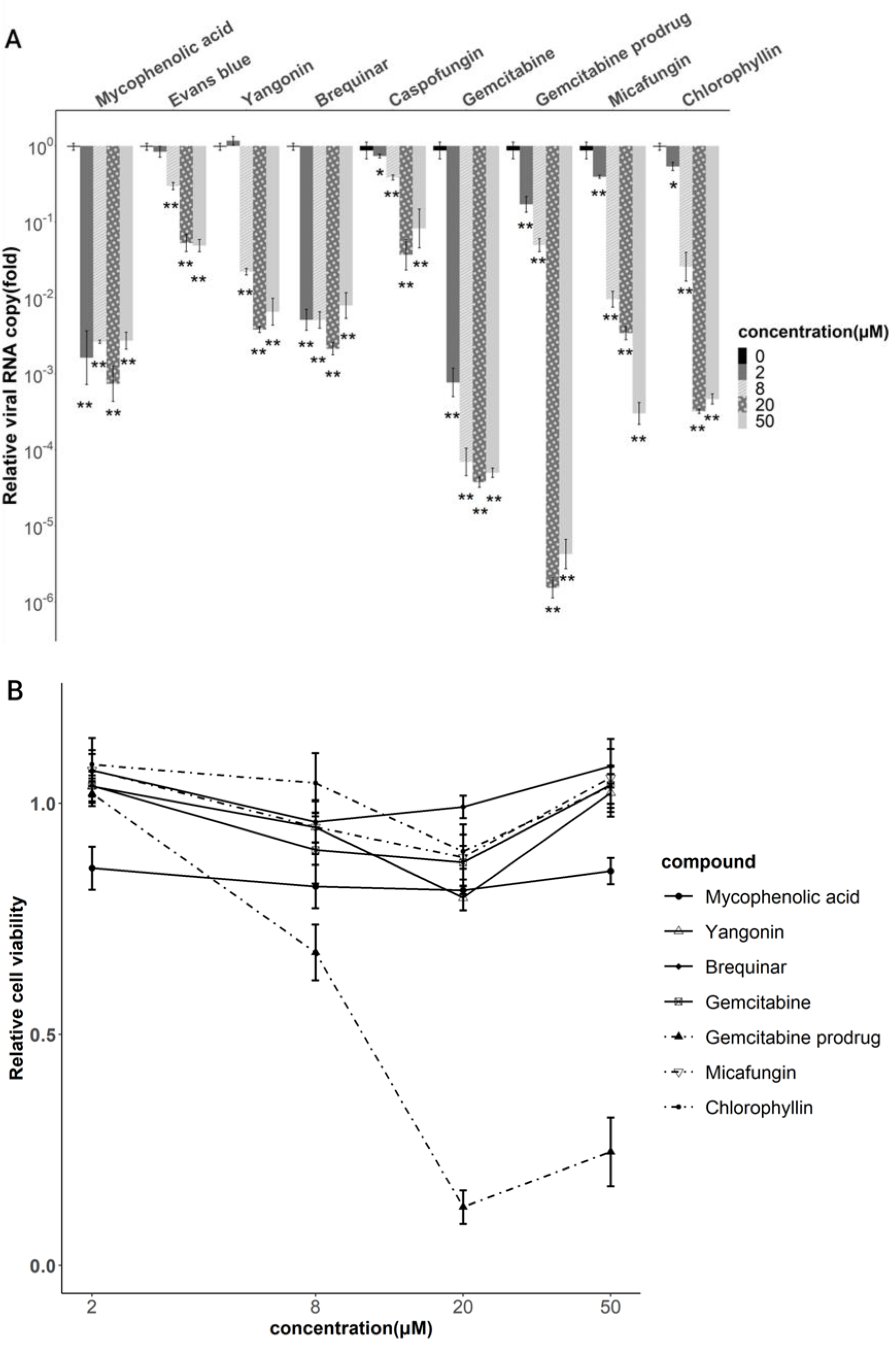
Validation of anti-PRV effect of hit compounds. (A) Dose-dependent viral replication inhibition of hit compounds. Vero cells treated with compounds and infected at MOI = 0.01 for 24 hours. The relative viral RNA copy was determined by RT-qPCR (n = 6, mean ± SE). Gemcitabine prodrug means 3’,5’-Di-O-benzoyl-2’-deoxy-2’,2’-difluorocytidine. Statistical analysis performed by package rstatix (version 0.7.0) wilcox_test function (*p < 0.05, **p < 0.01). (B) Cell cytotoxicity of hit compounds. The relative cell viability of Vero cells treated compounds for 24 hours by ATP assay (n = 6, mean ± SE).

### 3.3 Gemcitabine, Micafungin, and Yangonin inhibit PRV replication at the post-entry step

The viral infection cycle was divided into three steps; binding on the host cell membrane, entry into the host cell, and post-entry. Low-temperature treatment at 4 °C for 90 min enables the virus to bind the host cell membrane but inhibits its entry into the host cell. Using this mechanism, we performed a Binding / Entry / Post-entry assay to investigate in which step compounds inhibit PRV replication (Fig. 3A). Concentrations of each compound were set at that largely inhibited PRV replication in section 3.2 (gemcitabine, mycophenolic acid, brequinar = 2 µM, chlorophyllin, yangonin = 20 µM, micafungin = 50 µM). None of the compounds showed significant inhibition in the virus binding and entry steps (Fig. 3B). Gemcitabine, yangonin, and micafungin inhibited viral replication at the post-entry step at levels comparable to those observed in section 3.2 (Fig. 3B). Other compounds only reduced the relative viral copy number to 1.0 × 10^−1^.

**Fig 3.**
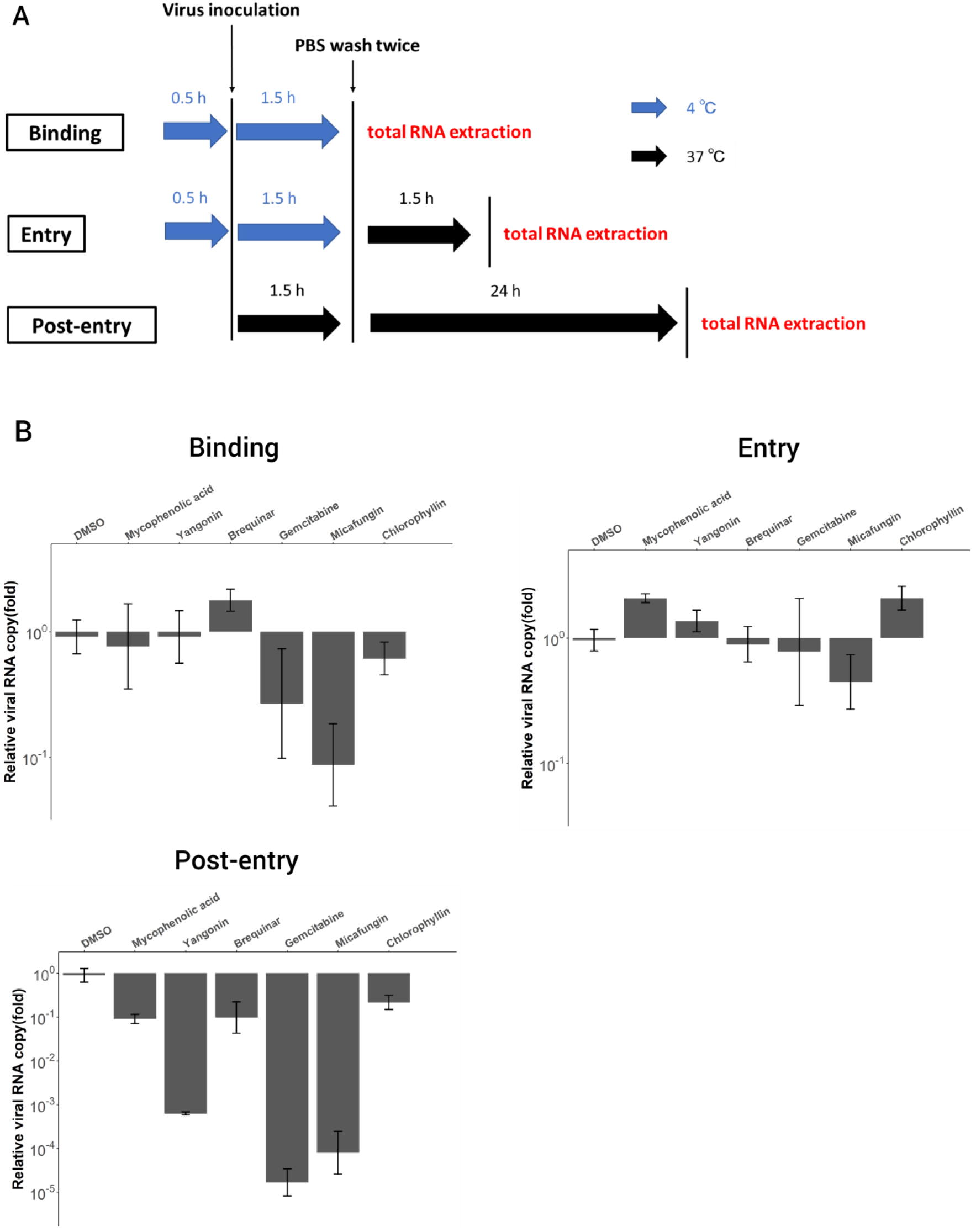
(A) The illustration of Binding / Entry / Post-entry assay protocol. (B) Results of RT-qPCR (n = 3, mean ± SE).

### 3.4 Micafungin inhibits another strain of PRVs at the post-entry step

Based on previous experiments, we focused on micafungin, which has a high repositioning advantage among the hit compounds. Then, to further expand the potential of micafungin as an anti-PRV agent, we tested its inhibitory effect on PRV69G, another strain of PRVs that we isolated in Indonesia. Micafungin significantly inhibited the replication of PRV69G and PRV50G at 50 µM (Fig. 4A and 4B).

**Fig 4.**
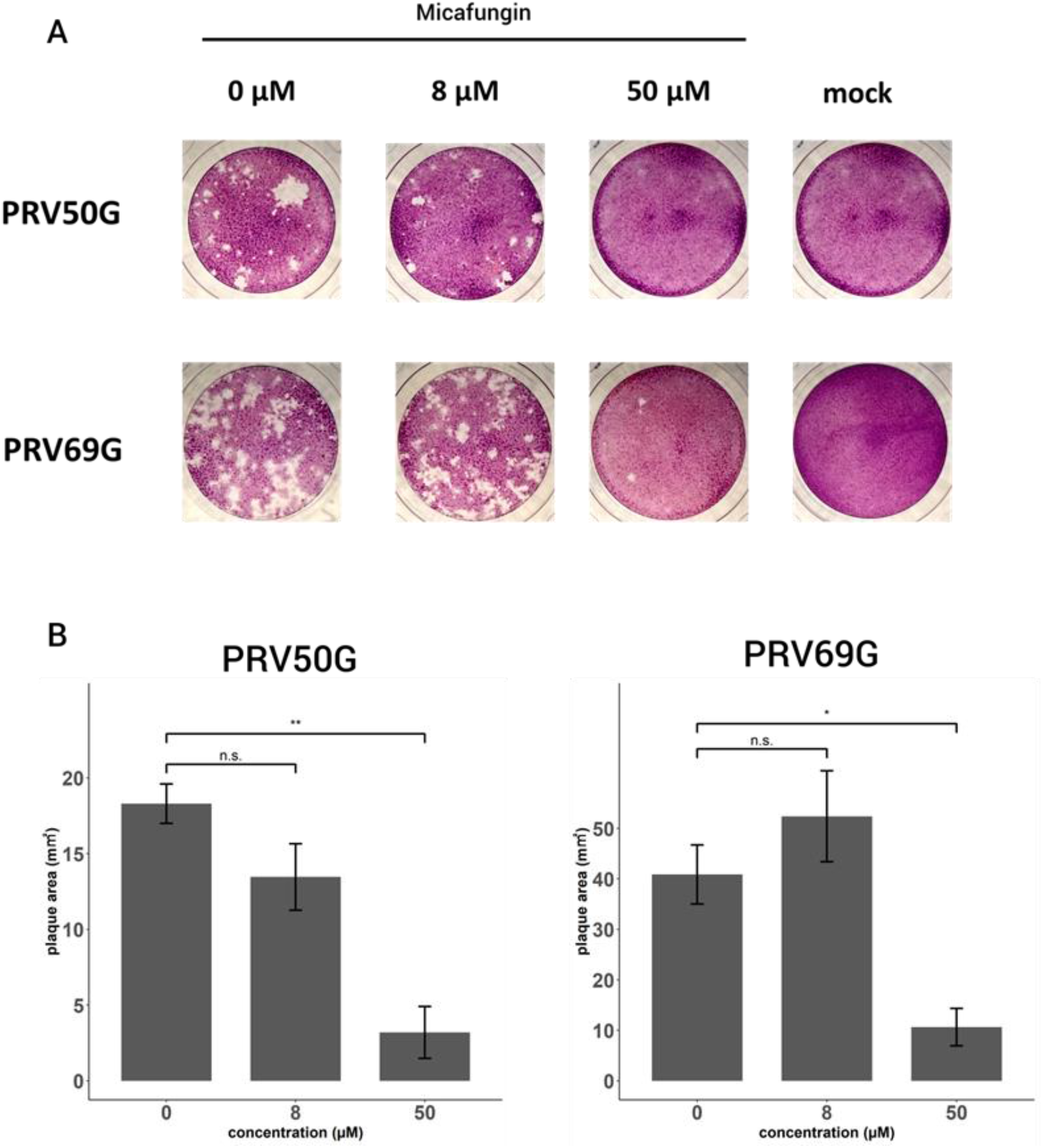
(A) Plaque assay of micafungin against PRV50G and PRV69G. (B) Area of plaques was calculated by ImageJ Fiji. Statistical significant was analyzed by Welch t-test (*p < 0.05, **p < 0.01, ns = not significant with 0 µM group).

## 4. Discussion

We identified eight hit compounds among 2,943 FDA-approved drugs that showed anti-PRV activity *in vitro*. The screening was performed in two steps: observation of viral cytopathic effect and MTT assay to measure cell viability (Fig. 1A). Dose-response analysis by RT-qPCR showed that six compounds, except evans blue and caspofungin, particularly inhibited PRV replication (Fig. 2A). Subsequent ATP assays revealed that the confirmed anti-PRV effect was not due to cell cytotoxicity (Fig. 2B). Hit compounds included mycophenolic acid, brequinar, and gemcitabine, which are already known for their inhibitory effect against rotavirus belonging to the same family *Reoviridae* (Fig. 1D) (Chen et al., 2020, 2019; Yin et al., 2016).

Because viruses use host nucleotides to replicate their genomes (Shatkin, 1969), depletion of host nucleotide pools leads to inhibit viral replication (Hoffmann et al., 2011). Mycophenolic acid and brequinar inhibit dehydrogenases required for *de novo* purine/pyrimidine biosynthesis (Chen et al., 1992.; Sintchak et al., 1996). Gemcitabine, a cytidine analogue, also inhibits pyrimidine biosynthesis (Lee et al., 2017). These three drugs inhibit multiple viral replications; hence they are known as broad-spectrum antiviral agents (Andersen et al., 2020). Furthermore, there are accumulating reports that the three drugs activate some interferon-stimulated genes (ISGs), an innate immune system of the host (Li et al., 2020; Luthra et al., 2018; Pan et al., 2012; Shin et al., 2018). Here we showed that these three drugs have inhibitory effects on PRV. The weakened effect of brequinar and mycophenolic acid was observed in the post-entry step (Fig. 3B). On the other hand, gemcitabine inhibits PRV replication at a high level in the post-entry step, which is thought to be due to its direct action on the viral polymerase (Shin et al., 2018). These results raise the possibility of repositioning gemcitabine against PRV, that currently widely used as an anticancer agent. In this study, gemcitabine already inhibits PRV replication at the point of 2 µM, which is consistent with previous results with rotavirus (Chen et al., 2020). In murine leukemia virus (Clouser et al., 2011), HIV-1 (Clouser et al., 2012), and human rhinovirus (Song et al., 2017), studies showed that antiviral activity of gemcitabine *in vivo* achieved at doses significantly lower than those given as an anticancer agent. However, because strong side effects of gemcitabine, such as myelosuppression, are considered more severe than symptoms of PRV infection at present, careful consideration should be given to repositioning (Abbruzzese et al., 1991).

Yangonin is an extract of the kava plant, and chlorophyllin sodium copper salt (CHL) is a semi-synthetic compound derived from the naturally-occurring green pigment chlorophyll and used as a food additive (Clouatre, 2004; Tumolo and Lanfer-Marquez, 2012). Antiviral effects on yangonin and CHL have been reported in several RNA viruses (Li et al., 2017; Liu et al., 2020a). However, the clinical use of yangonin has been restricted due to its hepatotoxicity (Clouatre, 2004). CHL has been reported to inhibit virus entry into host cells in enterovirus 71, coxsackievirus (Liu et al., 2020b). The inhibition mechanism could not be identified in this study.

Micafungin and caspofungin are echinocandins that specifically inhibit 1,3-β-glucan synthase, which synthesizes the fungal cell wall (Odds et al., 2003). Recently, the antiviral activity of micafungin and its analogues, caspofungin, and anidulafungin, have been reported in enterovirus 71 (Kim et al., 2016), chikungunya virus (Ho et al., 2018), dengue virus (Chen et al., 2021), zika virus (Lu et al., 2021), SARS-CoV-2 (Ku et al., 2020). The fact that these antifungals are already in use, and have known few side effects, is an advantage in repositioning (Cheng et al., 2022). Micafungin and caspofungin were identified as hit compounds in this screening, but micafungin was more effective in inhibiting PRV replication. Plaque assay revealed that micafungin inhibits another strain of PRVs, PRV69G. Phylogenetic analysis suggests that reassortment events occurred among various strains of PRVs (Takemae et al., 2018a). In mammalian orthoreovirus, frequent reassortment events have been reported in bats (Lelli et al., 2015; Naglič et al., 2018). The broad anti-PRV activity of micafungin should be effective against new reassortants that may emerge in the future.

Finally, we screened the FDA-approved drug library and identified antiviral drug candidates for PRV, a bat-borne emerging zoonotic virus. Three of the eight hit compounds were nucleic acid synthesis inhibitors, and two were antifungals. Micafungin is a drug with a powerful repositioning advantage; further *in vivo* experiments will expand its potential as an anti-PRV agent.

## Declaration of competing interest

The authors declare no competing interests.

## Notes

### Competing Interest Statement

The authors have declared no competing interest.

